# Epigenome-wide association study of asthma and wheeze in childhood and adolescence

**DOI:** 10.1101/135301

**Authors:** Ryan Arathimos, Matthew Suderman, Gemma C Sharp, Kimberley Burrows, Raquel Granell, Kate Tilling, Tom R Gaunt, John Henderson, Susan Ring, Rebecca Richmond, Caroline L Relton

**Author notes:** Corresponding author: Ryan Arathimos, MRC Integrative Epidemiology Unit, University of Bristol, Oakfield House, Bristol, BS8 2BN, UK.

## Abstract

Asthma heritability has only been partially explained by genetic variants and is known to be sensitive to environmental factors, implicating epigenetic modifications such as DNA methylation in its pathogenesis.

Using data collected in the Avon Longitudinal Study of Parents and Children (ALSPAC), we assessed associations of asthma and wheeze with DNA methylation at 7.5 years and 16.5 years, at over 450,000 CpG sites in DNA from the peripheral blood of approx. 1000 participants. We used Mendelian randomization (MR), a method of causal inference that uses genetic variants as instrumental variables, to infer the direction of association between DNA methylation and asthma.

We identified 302 CpGs associated with current asthma status (FDR-adjusted P-value <0.05) and 445 with current wheeze status at 7.5 years, with substantial overlap between the two. Genes annotated to the 302 associated CpGs were enriched for pathways related to movement of cellular/subcellular components, locomotion, interleukin-4 production and eosinophil migration. All associations attenuated when adjusted for eosinophil and neutrophil cell count estimates. At 16.5 years, two sites were associated with current asthma after adjustment for cell counts. The CpGs mapped to the *AP2A2* and *IL5RA* genes, with a -2.32 [95% CI -1.47,-3.18] and -2.49 [95% CI -1.56,-3.43] change in percentage methylation in asthma cases respectively. Two-sample bi-directional MR indicated a causal effect of asthma on DNA methylation at several CpG sites at 7.5 years. However, associations did not persist after adjustment for multiple testing. There was no evidence of a causal effect of asthma on DNA methylation at either of the two CpG sites at 16.5 years.

The majority of observed associations are driven by higher eosinophil cell counts in asthma cases, acting as an intermediate phenotype, with important implications for future studies of DNA methylation in atopic diseases.

## Introduction

Asthma continues to be a global burden on health care systems with 8.2% of the US population affected in 2009^(1)^ and 300 million affected worldwide ^(2)^. The prevalence of paediatric asthma in particular is on the rise along with modern lifestyle factors such as air pollutants and indoor contaminants and allergens thought to be influencing the increase^(3)^. Although asthma is known to be strongly heritable^(4)^, genome wide association studies (GWAS) have only been able to explain a fraction of that heritability^(5, 6)^.

As asthma development and progression are affected by environmental factors, a hypothesised role for epigenetic mechanisms has been proposed ^(7-9)^. There has been particular interest in a role for DNA methylation, one of the most widely studied epigenetic mechanisms which is known to be responsive to environmental exposures^(10)^. DNA methylation variation has previously been found to be associated with known asthma triggers, including air pollution ^(11, 12)^ and smoking^(13, 14)^. Whereas childhood asthma is often allergic in character and is more common in boys than girls^(15, 16)^, adult asthma is known to differ in that it is often more severe, non-atopic and leads to a more rapid decline in lung function^(16)^, indicating different biological mechanisms involved in the pathophysiology between asthma in childhood and asthma in late adolescence.

Wheeze is a symptom often observed as a precursor to children that develop asthma^(17, 18)^ and has been indicated to have important prognostic value in early detection of asthma^(19)^. Whereas wheeze and asthma are related traits, wheezing is a broader and more heterogeneous phenotype than asthma that encompasses a wider variety of respiratory conditions, that may include disorders such as Chronic Obstructive Pulmonary disorder (COPD), viral infection and bronchiolitis^(20)^. Associations of wheeze with DNA methylation may be expected to capture a wider variety of biological processes that underlie respiratory conditions.

Previous studies have reported an association between peripheral blood DNA methylation and IgE ^(21, 22)^, an important mediator of atopic or inflammatory diseases such as asthma. However, whether these associations are also present for the asthma phenotype has not been assessed. Studies that have examined DNA methylation associations with asthma have not specifically determined whether observed associations are due to cell type confounding between samples ^(23, 24)^. Eosinophils, a type of circulating granulocyte, are known to be increased in peripheral blood of some people with allergic conditions, such as certain subtypes of asthma ^(25, 26)^. Eosinophils are involved in the development of characteristic features of asthma, such as airway remodelling and hyperreponsiveness, heightened immune response and initiation of allergic airway inflammation ^(27, 28)^. The increase of eosinophils in some individuals with asthma may lead to “confounding” or contamination of whole-blood methylation data^(29, 30)^, as the distinct methylation patterns of eosinophils^(31)^ are over-represented in cases.

Associations observed between DNA methylation and disease outcomes, like all observational associations, are vulnerable to confounding, reverse causation and bias. Determining the direction of association is of critical importance to the biological interpretation of such findings^(32)^. Mendelian randomization (MR) is an approach that uses genetic variants as instrumental variables (IVs) and has been widely implemented to strengthen causal inference^(33, 34)^. Any association observed between the IVs and the disease outcome is most likely explained by an unbiased causal effect of the exposure on the outcome, given a certain set of prior assumptions^(34)^. MR can make use of the strong *cis* effects of genetic variants on DNA methylation. *Cis*-genetic variants robustly associated with site specific DNA methylation can be identified and used as causal anchors in such analyses ^(21, 35)^. This approach is particularly useful in the case of DNA methylation, where the causal direction of association between DNA methylation and the disease outcome may be difficult to establish using observational data alone. Furthermore, the application of MR is increasingly facilitated by the availability of GWAS data from large scale consortia as the methods can be applied using summary level statistics ^(36)^.

In the present study, we examine genome-wide DNA methylation in relation to asthma and wheeze in childhood and adolescence in a large contemporary UK-based cohort. We implement a variety of analytical strategies to gain insights into causal relationships of the DNA methylation variation observed.

## Methods

### Study population

The Avon Longitudinal Study of Parents and Children (ALSPAC) recruited 14,541 pregnant women resident in the former county of Avon, UK with expected dates of delivery 1st April 1991 to 31st December 1992. The initial number of pregnancies was 14,541, for which the mother enrolled in the ALSPAC study and had either returned at least one questionnaire or attended a “Children in Focus” clinic by 19/07/99. Of these initial pregnancies, there was a total 14,062 live births and 13,988 children who were alive at 1 year of age. The phases of enrolment are described in detail in the cohort profile paper ^(37)^. The study website contains details of all the data that is available through a searchable data dictionary http://www.bris.ac.uk/alspac/researchers/data-access/data-dictionary/.

Written informed consent has been obtained for all ALSPAC participants. Ethical approval granted from the ALSPAC Law and Ethics Committee and the local Research Ethics Committee in accordance with the guidelines of the Declaration of Helsinki.

#### Respiratory phenotypes

Estimates of current asthma/wheeze status and ever asthma/wheeze were derived from questionnaires completed by the mothers of the study children when they were 7.5 years-old and 16.5 years old respectively. Current asthma case status at 7.5 years of age was defined using a positive answer to the question “Has your child ever been diagnosed with asthma by a doctor?” and a positive answer to one of the following questions: (1) “Has your child had asthma in the past 12 months?” (2) “Has your child been on asthma medication the last 12 months?” or (3) “Has your study child had wheezing or whistling on the chest in the past 12months?”. Controls were children with negative answers to all the above questions. Current wheeze at 7.5 years was defined using answers to the question “Has your child had wheezing or whistling on the chest in the last 12 months?” and is therefore a broader definition of current asthma at that age. Current asthma case status at 16.5 years was defined using the same method, but from counterpart questions asked in the child-completed questionnaire. Ever asthma and ever wheeze at 16.5 years were defined using answers to the questions from questionnaires completed at 16.5 years of age by the mothers of the study participants. Answers to the questions “Has he/she [your study child] ever had asthma?” and “Has he/she ever had wheezing or whistling in the chest at any time in the past?” were used to define each measure respectively.

#### DNA methylation data

DNA methylation data for a sub-set of approximately 1000 mother-child pairs from ALSPAC is available as part of the ARIES (Accessible Resource for Integrated Epigenomics Studies) project^(38)^. ARIES participants were selected based on availability of DNA samples at three time-points for the offspring (neonatal, childhood and adolescence). Mothers included in ARIES were slightly older, more likely to have a non-manual occupation and less likely to have smoked throughout pregnancy when compared to the total ALSPAC study cohort^(38)^. DNA methylation was measured using the Illumina Infinium HumanMethylation450 BeadChip. DNA methylation at each CpG site is expressed as a beta-value (β) (ratio of methylated probe intensity to overall intensity representing 0 to 100% methylation at the CpG) based on the proportion of detected CpGs that were methylated at each site on the array^(39)^. Pre-processing of peripheral blood and cord blood samples was performed as previously described^(38)^. Details of quality control (QC) of the DNA methylation data can be found in the **Supplementary Material**.

#### Other variables

Proportions of cell types were estimated from DNA methylation data using the estimateCellCounts function in the minfi R package ^(40)^ which is based on the method developed by Houseman et al.^(41)^. This estimated the proportion of B cells, CD8 T cells, CD4 T cells, granulocytes, eosinophils, neutrophils, NK cells and monocytes at the 7.5 year methylation time-point and at the 16.5 year methylation time-point independently. Descriptive statistics of the derived cell counts can be seen in **Supplementary Table S1**. All epigenome-wide association study (EWAS) analyses were adjusted for three levels of cell-type adjustment. Model 1: unadjusted for cells, Model 2: basic cells (where eosinophils and neutrophils were grouped together as granulocytes) and Model 3: detailed cells (eosinophils and neutrophils assessed separately).

Atopic status was determined in a subset of ALSPAC children when they were approximately 7.5 years old by skin prick test response to a panel of up to 12 common allergens, including house dust mite, grass pollen, cat, egg, peanut, and mixed nuts. A positive response was defined as a mean weal diameter of >2 mm with an absent response to negative control solution, and atopy was defined as a positive response to one or more of house dust mite, grass pollen, and cat.

### Statistical analyses

We first examined cross-sectional associations of asthma status at both 7.5 years and 16.5 years with estimated cell counts. We used linear regression models in Stata 13 MP2 ^(42)^ to determine associations between cell types that were most likely to have a strong effect on results when adjusted for in EWAS models.

#### Epigenome-wide association studies

EWAS were carried out in R version 3.0.2. using the CpGassoc package^(43)^ to model either asthma/wheeze status as the exposure and methylation (beta-value) at either 7.5 years or 16.5 years of age as the outcome. Results were then corrected for multiple testing by controlling the expected proportion of false-positives among all discoveries (FDR-adjusted P-value < 0.05), and type I errors were controlled using Bonferroni adjustment. All models were adjusted for sex, maternal smoking during pregnancy, maternal age, parity, maternal education derived from maternal questionnaires and unknown confounders using surrogate variables (by use of maximum 10 surrogate variables obtained from the sva R package^(44)^), as shown in **Supplementary Table S2.** Probes that were known to contain SNPs or be of low quality^(45)^ were removed. The genomic inflation factor (Lambda λ) and quantile-quantile (Q-Q) plots were used to compare the genome-wide distribution of P-values with the expected null distribution.

We examined three models: Model 1 was adjusted offspring sex, maternal age, parity, smoking status, maternal education and for unknown confounders using surrogate variables. Model 2 adjusted additionally for basic cell counts (which grouped eosinophils and neutrophils under granulocytes) and Model 3 adjusted for covariates as above and detailed cell counts.

#### Sensitivity analyses

We performed a sensitivity analysis where individuals with extreme values of derived eosinophil cell counts at 7.5 years were removed using Tukey’s method of outlier removal ^(46, 47)^.This was done as some evidence suggest eosinophilic asthma represents a subtype of asthma within a wider spectrum of asthmatic disorders^(48, 49)^. We used both a stringent and a relaxed version of Tukeys method. For the stringent method potential outliers in the eosinophil cell count data were removed if their value was more than the upper quartile plus one and a half times the interquartile range (IQR). For the relaxed method outliers were removed if their value was more than the upper quartile plus three times the interquartile range (IQR). We did not remove values that were less than the lower quartile minus three times the interquartile range as having an eosinophil cell count of zero (or near-zero) is normal for most individuals that do not have an allergic disease. We also stratified the outlier removal by sex to determine whether outliers in eosinophil cell counts were more frequent in females or males.

#### Functional analysis

The Gene Ontology (GO)^(50)^ database in DAVID^(51)^ Bioinformatics Resources 6.8 (beta) was used to examine gene function in potential molecular, cellular and biological processes of loci observed to be associated with asthma at age 7.5 years (adjusted for basic cell counts). The Kyoto Encyclopedia of Genes and Genomes (KEGG) database^(52)^ in DAVID was used to explore whether genes were related to known pathways involved in asthma or eosinophil regulation.

#### Mendelian randomization

In order to investigate the causal relationships between methylation and asthma, a MR approach was applied. Briefly, MR is a method of investigating causal relationships by using genetic variants as instrumental variables (IVs)^(33)^. The main assumptions and strengths of the technique have been outlined in detail elsewhere^(53)^. A two-sample approach to MR was used which has the additional benefit of only requiring summary level data from an outcome GWAS and an exposure GWAS. Consequently, these analyses are able to take advantage of published GWAS summary statistics from large consortia. The strengths and limitations of this specific technique have been reviewed ^(54, 55)^. A bi-directional approach was used in order to assess both the potential causal effect of asthma on DNA methylation and the effect of DNA methylation on asthma. The TwoSampleMR package in R available as part of the MR-Base (http://www.mrbase.org) platform ^(36)^ was used for all MR analyses. We analysed the causal effect of the asthma SNPs on CpGs from the EWAS of asthma at 7.5 years adjusted for basic cell counts.

For assessing the causal effects of asthma on DNA methylation, asthma SNPs from the GWAS catalog ^(56)^ were used as IVs for asthma and methylation data from the ARIES dataset was used as our second sample (**Supplementary Figure S2**). Specifics on the selection and QC of SNPs from the GWAS catalog used as IVs can be found in the **Supplementary Material**. A combination of methods, including the inverse-variance weighted method, maximum likelihood method, weighted median method and MR-Egger regression, were used as sensitivity analyses. We tested for pleiotropy by assessing the MR-Egger regression intercept with funnel plots/scatter plots and for heterogeneity using Cochran’s Q statistic.

In the reverse direction, *cis*-SNPs robustly associated with methylation levels at the specific CpG sites were used as proxy IVs for DNA methylation^(57)^. For assessing the causal effect of DNA methylation on asthma we identified *cis*-SNPs associated with EWAS hits in ARIES and assessed the association of these *cis*-SNPs with asthma using summary data from the GABRIEL consortium GWAS of asthma^(6)^ as the second sample (**Supplementary Figure S3**). The Wald ratio method was used in the DNA methylation to asthma direction as only one *cis*-SNP was selected per CpG due to high Linkage Disequilibrium (LD) between neighbouring SNPs.

We calculated the effective number of independent tests for both the asthma to methylation and the methylation directions of the MR analysis. Adjustment for multiple testing is required when large numbers of tests are performed in an MR framework. However, CpGs are likely to be co-methylated if they are located in the same gene or close to each other on the chromosome. In order to take in to account potential correlation between nearby CpGs we calculated the effective number of independent tests for the 302 CpGs using matSpDlite^(58)^. Briefly, the method determines the number of independent tests using spectral decomposition of the correlation matrix of the tested exposures or outcomes. It calculates the equivalent number of independent variables in the correlation matrix, by examining the ratio of observed eigenvalue variance to its theoretical maximum. The method calculates a p-value at which type-I errors are kept at 5% based on the Veff process described by Li & Ji ^(59)^.

We performed power calculations using the mRnd Mendelian randomization power calculator by CNS genomics available online^(60)^. In the asthma to DNA methylation direction we had a maximum 6% power to detect a 3% change in methylation. In the DNA methylation to asthma direction we had a maximum 73% power to detect an OR of 1.1. Details of the power calculation can be found in **Supplementary Material**. Due to the low power to detect causal effects using Mendelian randomization, we present results for all CpGs from the EWAS and discuss in detail results from the top 20 CpGs based on lowest P-value in the EWAS only, as they demonstrate the most robust association with asthma status. Specific details of the MR process can be found in the **Supplementary Material.**

## Results

### Sample characteristics

Descriptive statistics of ARIES individuals included in each EWAS model are shown in **Table 1**. Individuals included as cases in each model were more likely to have a mother that smoked when compared to controls. Therefore, we adjusted for maternal smoking in all models as well as other potential confounders (maternal education, parity and maternal age).

**Table 1.** Characteristics of the individuals in each EWAS model.

| Methylation<br>(outcome) | Asthma/<br>wheeze<br>(exposure) | Status | N | Sex [Male]<br>N(%) | Maternal age<br>[months]<br>mean (SD) | Maternal<br>smoking<br>[Never] N(%) | Parity<br>[Nulliparous]<br>N(%) | Maternal<br>education<br>[O-level or<br>lower] N(%) |
| --- | --- | --- | --- | --- | --- | --- | --- | --- |
| 7.5 years | Current<br>asthma at<br>7.5 years | Cases | 149 | 88 (59%) | 369 (53.4) | 124 (83%) | 62 (42%) | 68 (46%) |
|  |  | Controls | 632 | 308 (49%) | 364 (51.7) | 560 (89%) | 286 (45%) | 306 (48%) |
|  | Current<br>wheeze at<br>7.5 years | Cases | 111 | 66 (60%) | 372 (55) | 94 (85%) | 43 (39%) | 520 (45%) |
|  |  | Controls | 742 | 366 (49%) | 361 (52) | 650 (88%) | 348 (47%) | 362 (49%) |
| 16.5 years | Ever asthma<br>at 16.5 years | Cases | 194 | 102 (53%) | 363 (53) | 163 (84%) | 97 (50%) | 87 (45%) |
|  |  | Controls | 554 | 264 (48%) | 364 (51) | 503 (91%) | 258 (47%) | 262 (47%) |
|  | Ever wheeze<br>at 18.5 years | Cases | 204 | 108 (53%) | 367 (49) | 173 (85%) | 102 (50%) | 82 (40%) |
|  |  | Controls | 554 | 264 (48%) | 363 (52) | 503 (91%) | 261 (47%) | 271 (49%) |
|  | Current<br>asthma at<br>16.5 years | Cases | 184 | 73 (40%) | 362 (51.6) | 150 (82%) | 89 (48%) | 89 (46%) |
|  |  | Controls | 427 | 200 (47%) | 364 (51.7) | 384 (90%) | 204 (48%) | 198 (46%) |

There was strong evidence of associations between current asthma and estimated cell counts, particularly in childhood (**Table 2**). Current asthma at 7.5 years demonstrated a strong positive association with estimates of derived B-cells and eosinophils, with a 0.026 [95%CI 0.018, 0.033] increase in the proportion of B-cells and a 0.011 [95%CI 0.005, 0.018] increase in the proportion of eosinophils in individuals with asthma respectively. There was a strong negative association with current asthma at the same age for neutrophils, with a 0.042 [95%CI -0.06, -0.022] decrease in proportion of neutrophils in individuals with asthma. Similarly, current asthma at 16.5 years demonstrated a positive association with estimates of derived eosinophils at 16.5 years with a 0.005 [95%CI 0.002, 0.008] increase in proportion, although with markedly less confidence in the estimate than the associations at 7.5 years. There was no evidence of an association between asthma at either 7.5 years or 16.5 years and any of the other cell types. A graphical depiction of the distributions of cell counts by asthma status at both 7.5 years and 16.5 years is shown in **Figure 1**. The same association with eosinophil cell counts was present for current wheeze at 7.5 years (**Supplementary Figure S4**)

**Table 2.** Associations of estimated cell counts at 7.5 years and 16.5 years with current asthma status at 7.5 years and 16.5 years respectively.

|  | Current asthma at 7.5 years |  | Current asthma at 16.5 years |  |
| --- | --- | --- | --- | --- |
|  | Change in proportion †<br>[95% CI] | P-value | Change in proportion †<br>[95% CI] | P-value |
| Monocytes | -0.002 [-0.005,0.004] | 0.695 | -0.003 [-0.007,0.004] | 0.534 |
| CD8T | 0.002 [-0.008,0.009] | 0.777 | -0.004 [-0.013,0.007] | 0.496 |
| CD4T | 0.009 [-0.002,0.018] | 0.069 | 0.009 [0,0.018] | 0.055 |
| B-cells | 0.011 [0.005,0.018] | <0.001 | 0.005 [-0.003,0.01] | 0.173 |
| NK cells | -0.004 [-0.008,0.002] | 0.159 | -0.004 [-0.01,0.004] | 0.306 |
| Neutrophils | -0.042 [-0.06,-0.022] | <0.001 | -0.009 [-0.029,0.011] | 0.382 |
| Eosinophils | 0.026 [0.018,0.033] | <0.001 | 0.005 [0.002,0.008] | 0.008 |

† change in proportion for individuals with asthma

**Figure 1.**
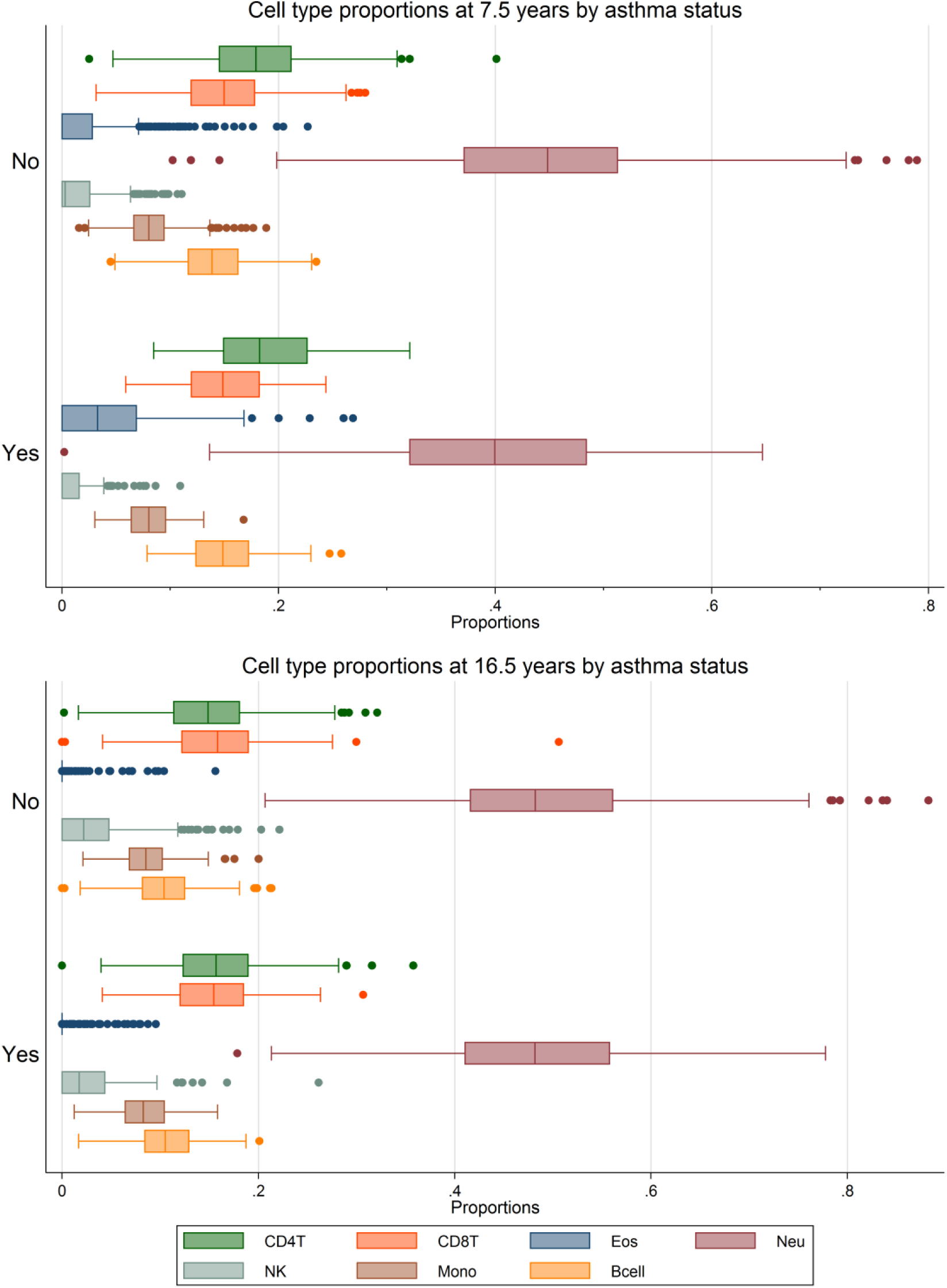
Box-plots of the estimated proportions of the 7 different cell types at 7.5 years and 16.5 years in individuals with asthma and individuals with no asthma at 7.5 years and 16.5 years respectively.

Associations of asthma at 7.5 years with cell counts stratified by atopic status indicate that increased eosinophils are evident in both atopic and non-atopic individuals with asthma. There is a 0.032 [95%CI 0.016, 0.047] difference in the proportion of eosinophils in atopic individuals with asthma (**Supplementary Table S4**), whereas for non-atopic individuals with asthma, the association is somewhat weaker with a 0.016 [95%CI 0.007,0.025] difference in the portion of eosinophils. For non-atopic individuals there is also evidence of an association of asthma with B-cells and neutrophils, with a 0.015 [95%CI 0.007,0.023] and -0.039 [95% CI -0.065, -0.014] change in proportion respectively.

### Epigenome-wide association analyses

We identified 411 CpG sites associated with current asthma status at 7.5 years (FDR-adjusted P-value <0.05) and 611 CpG sites with current wheeze status at 7.5 years. When adjusted for cell counts (eosinophils and neutrophils grouped together as granulocytes), 302 CpG sites remained associated with current asthma and 445 with current wheeze at 7.5 years. Adjusting for cell counts that included neutrophil and eosinophil counts separately (detailed cell counts) attenuated all associations (**Table** 3). We found evidence of genomic inflation in all unadjusted models that reduced progressively once adjustments for basic cell counts and detailed cell counts were made (**Supplementary Figure S6**). The top locus associated with current asthma at 7.5 years adjusted for cell counts that did not include eosinophils and neutrophils mapped to *ZFPM1* (cg04983687) with a -5.02 [95% CI -3.5, -6.5] difference in percentage methylation for individuals with asthma. A larger number of sites were associated with current wheeze at age 7.5 years (445 CpG sites), and the top locus mapped to *MFHAS1* (cg1207746) with a -4.60 [95%CI -3.33, -5.86] difference in percentage methylation for individuals with wheeze. There was a high degree of overlap between current asthma and current wheeze associated CpG sites (57.1% of 611 sites associated with current wheeze or 349 CpG sites) in the unadjusted model; 55.7% of 445 sites associated with current wheeze or 248 CpG sites in the adjusted model). Current asthma at 16.5 years was however associated with methylation at the same age at 2 CpG sites after adjustment for cell counts that included eosinophils and neutrophils. The CpGs were annotated to the genes *AP2A2* (cg17676835) and *IL5RA* (cg10159529) with a -2.3 [95% CI -1.47, -3.18] difference and -2.5 [95% CI -1.56, -3.43] difference in percentage methylation respectively (**Figure 2**). There were no associations observed between methylation at 16.5 years and ever asthma status at the same age. There were 32 CpGs associated with ever wheeze at 16.5 years once adjusted basic cells, but all associations attenuated once adjusted for cell counts that included eosinophils and neutrophils. Full results for all probes with a p-value <1e-5 can be found in **Supplementary Tables E1-E15**.

**Figure 2.**
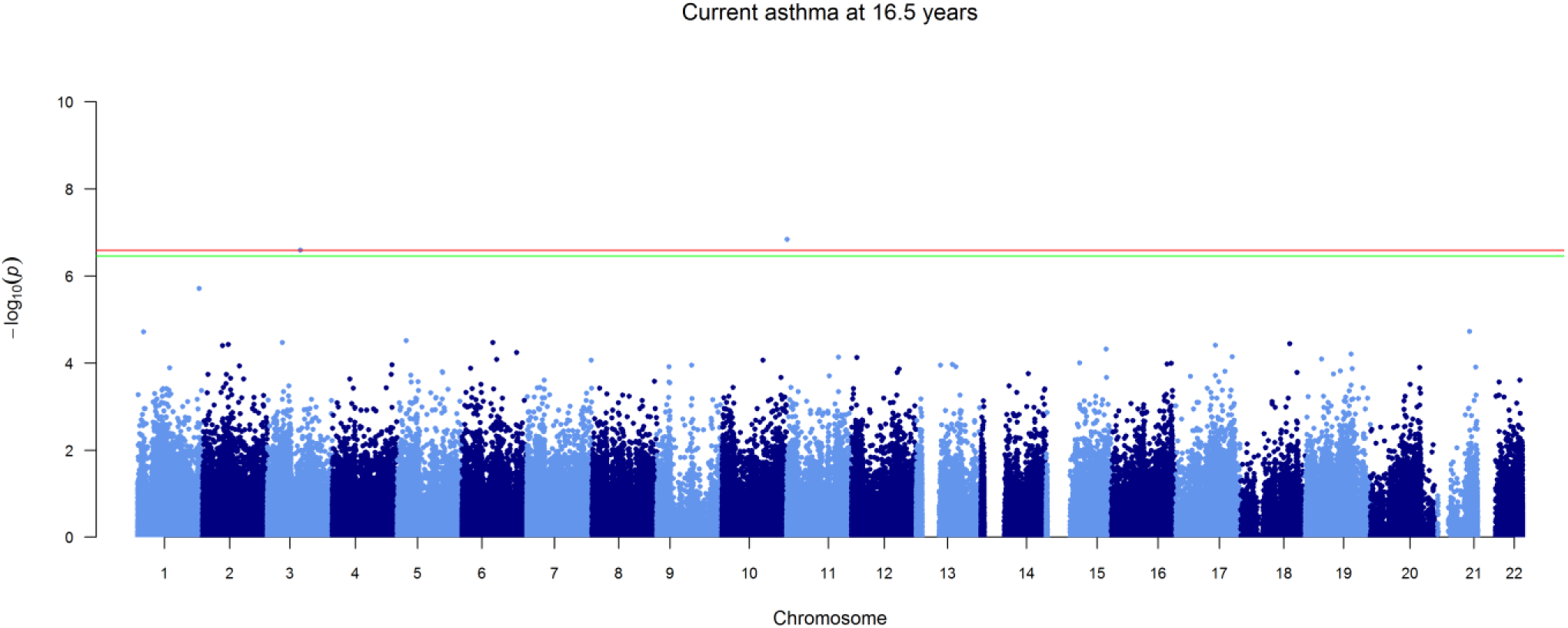
Manhattan plot of current asthma at 16.5 years adjusted for detailed cells. FDR significance cut-off line in green and Bonferroni cut-off line in red.

### Sensitivity analyses

In order to check whether associations of current asthma status at 7.5 years and cell counts were driven by outlier individuals with extreme levels of eosinophil cell counts, two outlier removal methods (Tukey’s method of outlier removal^(46)^, both stringent and relaxed) were applied to the eosinophil cell count data. **Supplementary Table S5** shows the effect of removal of outliers from the eosinophil cell counts. The association of asthma status with eosinophil cell counts remains even after stringent outlier removal. However, stratifying by sex shows that this appears to be a male phenomenon since there is no observed association in females (p > 0.05), as shown in **Supplementary Table S6**. The persistent association appears to be driven almost entirely by males. Based on this observation we conducted a sensitivity analysis where we restricted the EWAS sample to just females for current asthma at 7.5 years (N=385). There was one site associated with current asthma in the model unadjusted for cell counts (cg07612991), however once adjusted for basic cell counts, the association attenuated (**Supplementary Table E16, E17 & E18**). Power to detect an effect may have been greatly reduced in the model due to halving of the original sample size.

In order to ensure that adjustment for all 7 estimated cell counts was not having unintended effects due to multicollinearity between the estimates, we ran the same EWAS of current asthma at 7.5 years adjusted only for estimated eosinophils. However, we observed the same attenuation of associations as the model adjusted for all 7 estimated cell counts.

### Enrichment analysis/Functional analysis/Gene ontology

A look up in the GO ^(50)^ and KEGG ^(52)^ databases showed that the 192 genes annotated to the 302 hit CpGs from the EWAS of current asthma at 7.5 years adjusted for basic cells were located in pathways related to movement of cellular or subcellular components, locomotion, interleukin-4 production and eosinophil migration. The genes were enriched in the following KEGG-pathways: amino/nucleotide sugar metabolism and asthma (**Supplementary Table S7 and S8**)

## Mendelian randomization

### Asthma to DNA methylation

We found some evidence of a causal effect of asthma on DNA methylation at several CpG sites (**Supplementary Table E19**). One CpG was among the top 20 hits from the EWAS of current asthma at 7.5 years, cg12077460 annotated to gene *MFHAS1* (beta IVW=-0.022, p=0.013) shown in **Figure 3**. However, none of the causal effect estimates met the calculated significance threshold of p= 0.00034 calculated based on the number of effective independent tests (n=151).

In the two CpG sites associated with asthma at 16.5 years there was no evidence of a causal effect of asthma on DNA methylation, at either the *AP2A2* gene (cg17676835), or *IL5RA* (cg10159529) shown in **Supplementary Table E21**.

### DNA methylation to asthma

In the reverse direction, from DNA methylation to asthma, 31 out of 302 CpGs were found to have *cis*-SNPs that were available in the meta-analysis results of the GABRIEL asthma GWAS. We were therefore only able to test the causal association at these 31 CpGs. There was evidence for a causal association between one CpG (cg11938718) and asthma annotated to the *HPCAL1* gene Wald beta=-1.766 (p=0.0024) shown in **Supplementary Table E22**. The strength of the effect did not meet the significance threshold calculated based on the number of independent tests (n=31, adjusted p= 0.0025). The effect of the CpG on asthma was also in the reverse direction to the observational effect from the EWAS (shown in **Supplementary Table E2**).

Neither of the two CpG sites associated with asthma at 16.5 years had available *cis*-SNPs that could be used as IVs. We were therefore unable to test the causal association in the reverse direction.

## Discussion

We have found multiple differentially methylated CpG sites in peripheral blood of asthmatics compared to non-asthmatics in childhood. However, when adjustments for estimated cell counts were made, to take into consideration the differences in circulating eosinophils and neutrophils between cases and controls, all of the observed associations attenuated. Evidence of genomic inflation in all unadjusted models reduced progressively once adjustments for basic cell counts and detailed cell counts (where eosinophils and neutrophils were modelled separately) were made, indicating that confounding by cell type was present in the samples. This is in line with our expectations of eosinophils being the driving factor in the detected signals. As expected, we find a far weaker association of asthma at 16.5 years with eosinophil cell counts, as adult onset asthma is more often non-atopic than when compared to childhood asthma^(16)^. Overall we found markedly fewer associations at 16.5 years than at 7.5 years with respect to current asthma and DNA methylation at the respective ages. We did not find any differentially methylated CpG sites when comparing individuals with ever asthma compared to never having had asthma at 16.5 years of age. Adjustments for cell counts at 16.5 years of age did not have the same impact as adjustments at 7.5 years. This is consistent with the likelihood that childhood asthma does not affect DNA methylation directly in our sample, but does affect estimates of DNA methylation from a mixed cell population due to the association between asthma and the constitution of the cell population.

We found more associations between current wheeze and DNA methylation at 7.5 years (611 hit CpG sites) than for current asthma (411 hit CpG sites) at the same age. However we observe the same attenuation in effects when adjusting for cell counts that include eosinophils in the current wheeze model. In the unadjusted model of current wheeze the CpG site with the strong association (cg12077460) mapped to the *MFHAS1* gene, whereas the top association in current asthma mapped to *ZFPM1* (cg04983687).

In the EWAS of current asthma at 16.5 years, the two CpGs that remained significant after adjustment for detailed cells mapped to the *AP2A2* and *IL5RA* genes. In the EWAS of current asthma at 7.5 years, cg17676835 (*AP2A2*) was not associated with current asthma at 7.5 years when adjusted for basic cells, whereas cg10159529 (*IL5RA*) was strongly associated in the same direction (p=0.0001) but attenuated when adjusted for detailed cells (p=0.304).

Several differentially methylated CpG sites in the EWAS of current asthma at 7.5 years were located in genes that have known links to both eosinophil functions and allergic sensitisation, such as cg27469152 in 3’UTR region of the *EPX* gene. The gene encodes eosinophil peroxidase, an enzyme expressed in eosinophils and released at sites of parasitic infection or allergic stimulation ^(61, 62)^. Both current asthma at 16.5 years and ever wheeze at 16.5 years were negatively associated with methylation levels at *IL5RA*, a gene encoding alpha subunit of the interleukin 5 (IL5) receptor.

Decreased expression of IL5 membrane receptor on eosinophils has been previously linked to allergen stimulation and increased IL5 protein levels in the airways ^(63)^.The fact that the association remains significant after adjustment for detailed cell counts in the model of current asthma at 16.5 years may indicate incomplete adjustment for eosinophils. Genetic polymorphisms of the *AP2A2* gene have also previously found to be associated with bronchitis and chronic obstructive pulmonary disease (COPD)^(64)^.

### Cell type heterogeneity

The increase of eosinophils in some individuals with asthma presents challenges when assessing whole blood DNA methylation ^(29, 30)^, as the distinct methylation patterns of eosinophils^(31)^ are over-represented in cases. Correction for white blood cell heterogeneity can be implemented, however, doing so adjusts for known characteristics of the disease outcome (asthma) therefore making it difficult to decipher methylation-related differences that are truly linked to the disease outcome.

Adjusting for cell type heterogeneity that may exist between different samples in an EWAS is of critical importance^(29)^ and multiple different methods have been devised to address the issue, which may be considered as a form of confounding^(65)^, although this categorisation is dependent on the hypothesis being tested. However, when a specific cell type forms an intermediate phenotype with involvement in the pathogenesis and characteristic features of a disease, then the unadjusted signals may be of biological interest. Raised eosinophil levels have been suggested to be an intermediate phenotype or feature of asthma ^(66)^. In the case of asthma, detected CpG signal from eosinophils may tag potentially pathogenic genes that might be causal to the asthma phenotype, when expressed differentially in blood due to higher numbers of eosinophils. By adjusting for estimated eosinophil counts, that signal is attenuated but may still have biological relevance.

### Causal analyses

Methods of causal inference such as MR help to define direction of causation in detected associations. Our MR analysis suggested a causal effect of asthma on DNA methylation at several CpG sites associated with asthma at 7.5 years. However, none of the effects survived multiple testing, even after adjusting the p-value threshold to take in to account correlation between the outcomes. Similarly, in the reverse direction, from DNA methylation to asthma there was no robust evidence for causal effects. We used a combination of complementary methods to ascertain these causal effects in MR, including fixed effect meta-analysis of Wald ratios, the inverse-variance weighted (IVW) method and MR Egger regression. The two-sample MR analysis was underpowered due to small sample size used to determine the SNP-methylation association, comprising just over 1000 individuals (from the ARIES-ALSPAC sample). Future work should aim to increase power by increasing the sample size in the dataset.

Based on these results we also attempted a two-sample MR analysis of eosinophils and DNA methylation using previously published SNPs that have shown strong association with eosinophil cell counts in peripheral blood ^(67)^. However the derived IVs from the published GWAS results were too weak to detect an effect in our sample. Future studies may identify further sequence variants that associate with eosinophil counts and provide more powerful IVs for MR analysis.

The genetic instruments used in the asthma to DNA methylation direction of the MR analysis were relatively weak in comparison to other IVs used in previous MR studies. Asthma GWAS from which the IVs were sourced have identified SNPs that explain very little of the population variance in asthma in comparison to other traits^(6, 68)^. Furthermore, our strict LD R^2^ cut-off threshold of <0.1 reduced the total amount of SNPs used as IVs to just six. In the reverse direction, from DNA methylation to asthma, we included only one *cis*-SNP for each CpG tested since strong LD was observed between neighbouring SNPs that forced the exclusion of many *cis*-SNPs that would have otherwise been combined to form stronger IVs.

### Comparisons with previous literature

A previously published EWAS of total serum immunoglobulin E (IgE) identified 36 CpG sites associated with differential levels of IgE^(21)^. IgE is a marker of allergic asthma and as such would be expected to be implicated in the biological pathway leading to allergic or inflammatory disease^(69)^. Whereas IgE is a robust biomarker of inflammatory conditions, it may not accurately proxy asthma as it encompasses a wide variety of atopic or inflammatory conditions. Of the 36 identified CpGs in this previous study, 35 were analysed in our EWAS, with 15 associated with current asthma at 7.5 years (FDR <0.05), and 13 remaining associated after adjustment for basic cells (FDR<0.05). Comparisons of results can be seen in **Supplementary Table S11**. This corroboration of findings suggests that eosinophil specific methylation patterns may be directly involved in asthma pathogenesis via raised IgE.

A recent EWAS of IgE in Hispanic children^(22)^ reported associations with DNA methylation in CpGs annotated to the *ZFPM1 (cg04983687, cg08940169)* and *ACOT7* (cg09249800, cg21220721, cg11699125) genes. We observe similar strong associations with CpG sites annotated to the same genes in the unadjusted EWAS of current asthma at 7.5 years (**Supplementary Table E1 and E2**).

However, these associations attenuate following adjustment for eosinophils and neutrophils, unlike the previously reported results. Furthermore, we have excluded all CpGs except cg04983687 (*ZFPM1*) from the results of our analyses as they have previously been indicated to be low-quality probes due to the effects of repeats, SNPs, INDELs, and reduced genome complexity^(45)^. Differences between results may indicate residual confounding due to eosinophils in previous results or lack of power in our own analyses, due to a lower number of asthma cases compared to controls relative to the previous study. Additionally, we have used exclusively methylation-estimated cell counts to adjust for cell mixture as opposed to cell sorted estimates of the previous study, leading to potentially imprecise adjustments in our own analyses. In a sensitivity analysis we generated estimated cell counts using the Houseman algorithm and the 200 reference CpGs supplied in the supplementary material of the previous study^(22)^. Correlations between the estimated cell counts using the estimated reference CpGs and our own Houseman derived estimates were relatively low. For eosinophils the correlation was 0.77, for neutrophils 0.96, for NK cells 0.57, for CD4T cells 0.75, for B-cells 0.71, for CD8T cells 0.13 and monocytes 0.76. As a sensitivity analysis using these estimated cell counts we reran the EWAS of at 7.5 years, however there were no associations between DNA methylation and asthma (FDR<0.05).

A previously published EWAS of asthma that did not adjust for eosinophil cell counts found similar associations to our own unadjusted results^(23)^. The EWAS focused on children with atopic asthma living in the inner city with an African-American, Hispanic or Caribbean background. Although the study focused on identifying differentially methylated regions (DMRs) instead of single sites, many of the resulting associations mapped to genes such as *ACOT7* and *IL13* that were also identified in our own unadjusted analyses. Similarly, a study of asthma using methylation data from nasal epithelium observed associations between asthma and DNA methylation but did not specifically adjust for eosinophil cell counts^(24)^. The authors in both of these studies do not appear to have considered confounding difference in cell mixtures between study cases and controls.

### Strengths and limitations

Strengths of the study include the ability to compare two different time-points in the same cohort of individuals, with methylation from childhood and adolescence, forming a series of cross-sectional analyses. Our use of complementary approaches to answering each question, which includes the functional analysis, sensitivity analyses and bi-directional MR (with multiple different testing methods applied) gives greater strength to our conclusions.

Limitations include the low coverage of the HumanMethylation450 Beadchip array, including less than 2% of all CpG sites^(70)^. Combined with our various probe exclusions, this brings the overall coverage down to just a fraction of the full methylome. Our analyses rely on the use of methylation data generated from peripheral blood leukocytes (PBLs), which may not be the most appropriate tissue in which to study associations with asthma. Cells from other tissues may be more relevant, such as airway epithelial cells, buccal cells or cells from bronchoalveolar lavage. In addition, our cell counts are estimated from the methylation data using a reference dataset instead of being directly sorted and counted in the original blood-samples. Whereas the method has been previously validated^(71)^, and previous EWAS analyses have found strong correlations between estimated and cell sorted measures^(22)^, it may not perform well in predicting proportions of rarer cell types in the blood, such as eosinophils, as they may be subject to less favourable signal-to-noise ratios than high abundance cell types. However, we attempted to counter the potentially poor estimation of cell counts by using surrogate variables in our EWAS analysis, a reference free method of adjusting for unknown confounding factors. We did not assess to what extent our detected associations were explained by genetic variants, such as SNPs.

Finally, measurement error due to misreporting of asthma symptoms is another limitation. Since our outcomes are derived from self-report questionnaires from either the mothers of the study children or the study children themselves, they are subject to misreporting of symptoms or previous asthma diagnosis. Any misreporting of asthma/wheeze symptoms is likely to bias results towards the null.

However, a validation study linking childhood asthma self-reports in ALSPAC using linked electronic patient records from GP practices shows that parental reports of a doctor’s diagnosis agree well with GP-recorded diagnosis^(72)^. Mendelian randomization is robust to measurement error in the exposure due to the use of genetic variants but may be subject to other biases in exposure and outcome^(73)^.

## Conclusion

We have shown multiple significant associations between asthma status and peripheral blood DNA methylation in both childhood and adolescence. There is strong evidence to suggest that the observed associations are driven to a large extent by higher eosinophil counts in asthmatics in childhood. We have shown that the use of whole-blood 450K data in EWAS of allergic diseases, such as asthma, where eosinophils may be expected to play a contaminating or confounding role in disease cases, makes it difficult to disentangle true associations of the disease with methylation from associations that may be driven by the overrepresentation of eosinophils. Whereas further work is needed to devise methods of elucidating true associations from those that are due to cell type confounding, this does not fully preclude the biological relevance of the observed differential methylation as raised eosinophils are a constituent part of the asthma phenotype.

## Acknowledgements

We are extremely grateful to all the families who took part in this study, the midwives for their help in recruiting them, and the whole ALSPAC team, which includes interviewers, computer and laboratory technicians, clerical workers, research scientists, volunteers, managers, receptionists and nurses. The UK Medical Research Council and Wellcome (Grant ref: 102215/2/13/2) and the University of Bristol provide core support for ALSPAC. This publication is the work of the authors and Mr Ryan Arathimos and will serve as guarantors for the contents of this paper. This work was supported by the MRC Integrative Epidemiology Unit and the University of Bristol (MC_UU_12013_2, MC_UU_12013_5 and MC_UU_12013_8). The Accessible Resource for Integrated Epigenomics Studies (ARIES) was funded by the UK Biotechnology and Biological Sciences Research Council (BB/I025751/1 and BB/I025263/1). GWAS data were generated by Sample Logistics and Genotyping Facilities at the Wellcome Trust Sanger Institute and LabCorp (Laboratory Corporation of America) using support from 23andMe. The funders had no role in study design, data collection and analysis, decision to publish, or preparation of the manuscript.

We are grateful to Dr Philip Haycock for his work in curating GWAS data as part of MR-base. We are also grateful to Mr Sam Abbott for his help with ggplot and Mr Ryan Langdon for his help in implementing Mendelian randomization for eosinophils.

**Figure 3.**
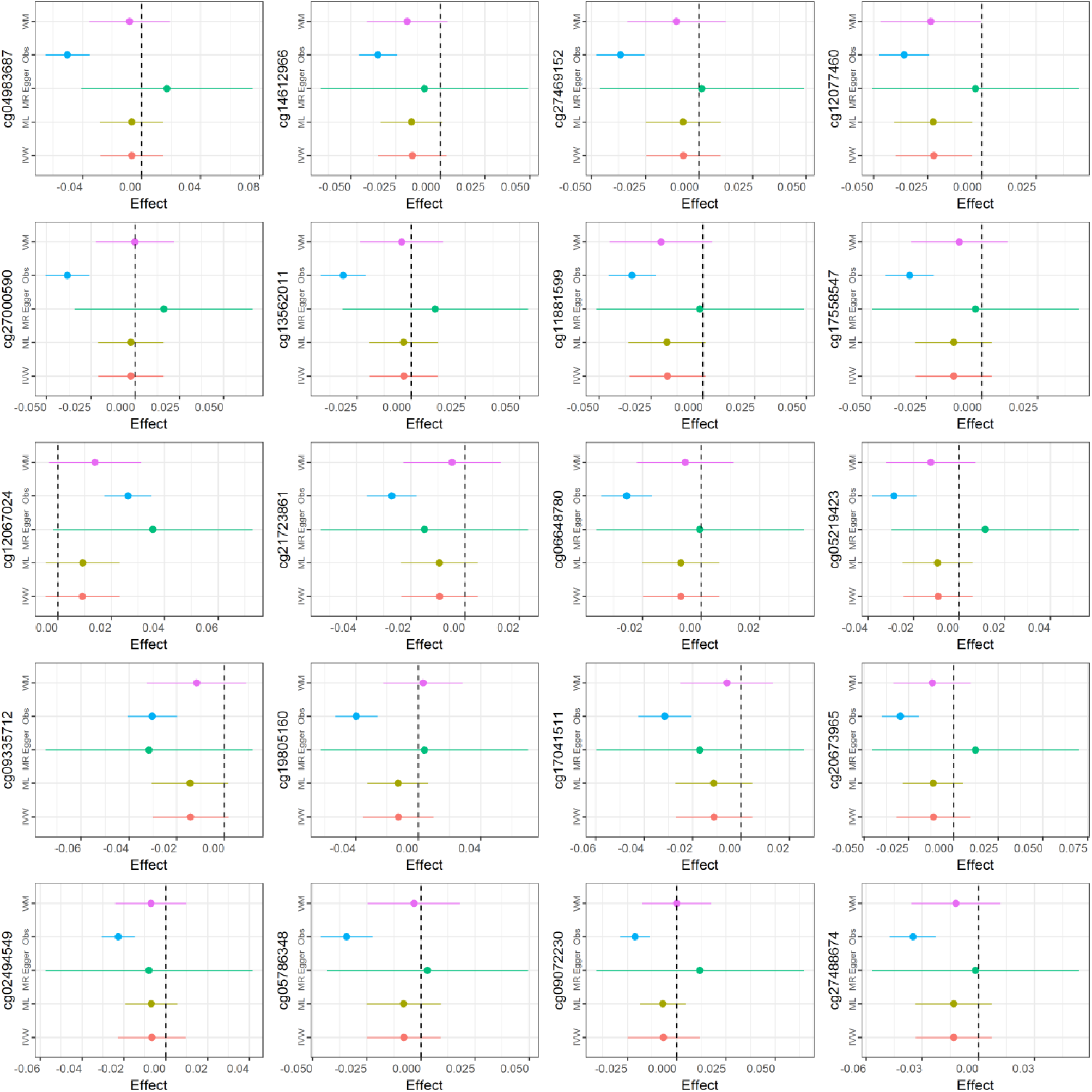
Causal effect of asthma on DNA methylation at each of top 20 CpGs from EWAS of asthma at 7.5 years and methylation at 7.5 years using two-sample Mendelian randomization approach. IVW= Inverse variance-weighted, ML= Maximum Likelihood, Obs=Observational effect from EWAS, WM= Weighted Median, MR Egger= MR Egger regression.

**Table 3.** Results of epigenome-wide association studies (EWAS) of methylation (outcome) and current asthma at 7.5 years, current wheeze at 7.5 years, current asthma at 16.5 years, ever asthma at 16.5 years and ever wheeze at 18.5 years (exposures). All models are adjusted for confounders and technical covariates. Stepwise adjustment for estimated cell counts were made (where basic cells include CD8T, CD4T, NK, monocytes, B-cells and granulocytes, and detailed cells include eosinophils and neutrophils instead of granulocytes). FDR-adjusted P-value and Bonferroni hits use a cut-off of <0.05. Lambda based on p-values of all probes without exclusions.

| Methylation (outcome) | Asthma/wheeze (exposure) | N [cases/controls] | Cell adjustment | FDR hits [Bonferroni] | lambda |
| --- | --- | --- | --- | --- | --- |
| 7.5 years | Current asthma at 7.5years | 781 [149/632] | No Cells | 411 [67] | 1.08 |
|  |  |  | Basic Cells | 302 [43] | 1.01 |
|  |  |  | Detailed Cells | 0 [0] | 0.97 |
|  | Wheeze at 7.5years | 853 [111/742] | No Cells | 611 [105] | 1.14 |
|  |  |  | Basic Cells | 445 [78] | 1.08 |
|  |  |  | Detailed Cells | 0 [0] | 1.00 |
| 16.5 years | Ever asthma at 16.5years | 748 [194/554] | No Cells | 0 [0] | 1.09 |
|  |  |  | Basic Cells | 0 [0] | 1.05 |
|  |  |  | Detailed Cells | 0 [0] | 1.01 |
|  | Ever wheeze at 16.5 years | 758 [204/554] | No Cells | 48 [9] | 1.02 |
|  |  |  | Basic Cells | 32 [9] | 1.05 |
|  |  |  | Detailed Cells | 0 [0] | 1.01 |
|  | Current asthma at 16.5 years | 611 [184/427] | No Cells | 2 [2] | 1.01 |
|  |  |  | Basic Cells | 2 [2] | 1.05 |
|  |  |  | Detailed Cells | 2 [1] | 0.99 |

